# Endothelial Caspase-9 Promotes Glial Changes, Inflammation, and Contrast Sensitivity Decline in Retinal Vascular Injury

**DOI:** 10.1101/2022.03.30.486450

**Authors:** Crystal Colón Ortiz, Albertine M. Neal, Maria I. Avrutsky, Monica Choi, Jade Smart, Jacqueline Lawson, Carol M. Troy

**Affiliations:** Department of Pathology & Cell Biology; Vagelos College of Physicians and Surgeons, Columbia University, New York, NY, 10032, USA; Barnard College, Columbia University, New York, NY, 10032, USA; Department of Neurology; Vagelos College of Physicians and Surgeons, Columbia University, New York, NY 10032, USA; The Taub Institute for Research on Alzheimer’s Disease and the Aging Brain; Vagelos College of Physicians and Surgeons, Columbia University, New York, NY 10032, USA

## Abstract

Retinal glial cells— microglia, astrocytes, and Müller glia—provide homeostatic support, regulate vascular blood flow, and react to injury by releasing inflammatory cytokines. Glial reactivity has been shown to be relevant for retinal vascular pathology and neuronal death. Non-apoptotic expression of endothelial caspase-9 (EC Casp9) was recently identified as a key mediator of retinal edema, hypoxic-ischemic injury, and neurodegeneration in retinal vein occlusion (RVO). In the current study we aimed to determine the glial responses that are modulated by EC Casp9 as a means to identify relevant neuro-immune mechanisms for the development of retinal edema and neurodegeneration. To this end we used a mouse model of RVO and a tamoxifen inducible EC Casp9 KO mouse line. We show that EC Casp9 leads to an increase in reactive microglia and to macrogliosis in a time-dependent manner. RVO induced an EC Casp9 dependent astroglial caspase-6 and cleavage of GFAP. Cytokine array analysis revealed that RVO increases expression of inflammatory cytokines out of which CX3CL1, IGF-1, IL-4, LIX, IL-1α, M-CSF, TNF-α, IL-1β, IL-10, and VEGF-A, were regulated by EC Casp9. Moreover, we found that EC Casp9 deletion resulted in protection from contrast sensitivity decline one day post-RVO. These results demonstrate that caspase-9 in hypoxic endothelial cells regulates retinal inflammatory signaling in microglia, astrocytes and Müller cells and changes in visual function.

## INTRODUCTION

The retina has three types of resident glial cells: microglia and macroglia (Müller glia, and astrocytes). Microglia are located in the synapse layers of the retina, the inner plexiform (IPL) and outer plexiform (OPL) layers, and the vasculature (Silverman & Wong, 2018). In close contact with the vasculature are the end feet of Müller glia and astrocytes in the retinal nerve fiber (NFL) and retinal ganglion layers (RGL). These glia form part of the blood retinal barrier (BRB) and control blood flow and vascular tone (Newman, 2015). Glial roles encompass regulating developmental functions, providing neuronal and vascular support through the release of neurotrophic factors and regulating immune responses and surveilling the retinal neuronal environment (Paisley & Kay, 2021; Reichenbach & Bringmann, 2020; Silverman & Wong, 2018). During retinal disease, glial cells undergo changes in morphology, become reactive, and release inflammatory cytokines that lead to neuronal dysfunction, recruitment of leukocytes, and breakdown of the BRB (de Hoz et al., 2016; Fletcher et al., 2008; Reichenbach & Bringmann, 2020; Silverman & Wong, 2018). Understanding the active interaction and contribution of glial cells to the dysregulation of the neuronal micro-environment and vasculature requires the characterization of the response of glial cells and thus their potential role in mediating neurodegeneration in retinal vascular disease.

Retinal vascular diseases such as retinopathy of prematurity (ROP), age-related macular degeneration (AMD), diabetic retinopathy (DR) and retinal vein occlusion (RVO), are major causes of vision loss that affect the quality of life (Brand, 2012). Retinal vascular disorders are characterized by impairment of the retinal vasculature through aberrant neovascularization and decreased blood flow that results in breakdown of the BRB, edema, inflammation, retinal neuronal atrophy, and visual dysfunction (Campochiaro, 2015). Vascular damage in the form of hypoxic-ischemic injury, such as is in RVO, is known to instigate a wide range of inflammatory responses. Many studies assessing vitreous samples of RVO patients have identified increased levels of pro-inflammatory cytokines and chemokines (Ehlken et al., 2015; Feng, Zhao, Zhang, Ma, & Jiang, 2013; Koss et al., 2012; Noma, Mimura, & Shimada, 2014; Shchuko et al., 2015). Similar results are observed in the mouse model of RVO (Fuma et al., 2017; Jovanovic, Liu, Kokona, Zinkernagel, & Ebneter, 2020). The presence of inflammatory mediators in patients with RVO is correlated with increased foveal thickness and hypoxic-ischemic injury (Noma et al., 2014), underlying the relevance of understanding the source of these cytokines in the injured retina. Current lines of treatment for RVO include anti-angiogenic and anti-inflammatory drugs. The first line of treatment targets vascular endothelial growth factor (VEGF) which acts on endothelial cells and promotes neovascularization that leads to edema (Le, 2017). Corticosteroid implants target inflammation and while they help to resolve macular edema they can also lead to cataract formation and increase of intraocular pressure (Arrigo & Bandello, 2021). Therefore, there is a need to better understand the inflammatory pathways in RVO to develop more specific therapeutics that simultaneously target neuroinflammation, neurodegeneration and retinal edema.

RVO leads to activation of neuroinflammatory responses in glial cells, including proliferation, migration and upregulation of pro-inflammatory responses that contribute to neurodegeneration. Microglial cells react to RVO with an increase in microglial cell number (Ebneter, Kokona, Schneider, & Zinkernagel, 2017) and participate in retinal ganglion cell (RGC) death (Jovanovic et al., 2020). Müller glia on the other hand, are known to present with increased markers of gliosis such as the intermediate filament, glial fibrillary acidic protein (GFAP), and redistribution and downregulation of the potassium channel Kir4.1 to perivascular areas (Koferl et al., 2014; Rehak et al., 2009). The astrocytic response and its pathological contribution to RVO, compared to microglia and Müller glia, has been understudied. Assessment of the level of GFAP, a marker of astrogliosis (Eng & Ghirnikar, 1994), in RVO has shown it to be either increased (McAllister et al., 2018), or unchanged (Koferl et al., 2014; Rehak et al., 2009). Increase of GFAP implicates changes in macroglial morphology and hypertrophy, which interferes with their normal function, although it is hypothesized that it could also be neuroprotective (Yang & Wang, 2015).

Our previous studies demonstrated that caspase 9, an initiator caspase, mediates neurodegeneration in models of cerebral ischemia (Akpan et al., 2011) and RVO (Avrutsky et al., 2020). Caspases are a group of cysteine dependent aspartate directed proteases that can activate cell death and inflammatory signaling. Over recent years their roles in the brain have been further characterized as relevant promoters of synaptic pruning, axon guidance, differentiation and in glial cells as mediators of inflammatory signaling and filament aggregation (Espinosa-Oliva, Garcia-Revilla, Alonso-Bellido, & Burguillos, 2019). Using a mouse model of RVO, we found non-apoptotic activation of EC Casp9. Pharmacological inhibition and genetic knock-out of EC Casp9 rescued neuronal death, BRB breakdown, edema, and potential modulation of inflammation (Avrutsky et al., 2020). However, whether EC Casp9 regulates the response of the resident glial cells to RVO remained to be answered. Understanding which resident glial cells and inflammatory cytokines are regulated by EC Casp9 mediated neurovascular injury will uncover pathways that are relevant for retinal edema and vision decline in RVO.

Here, we aimed to characterize the inflammatory signaling in RVO in the context of EC Casp9 deficiency. We found that EC Casp9 contributed to microglial and macroglial changes in a time-dependent manner. Additionally, we discovered that EC Casp9 modulated the levels of several cytokines and that its deletion rescued visual function decline in contrast sensitivity.

## MATERIALS AND METHODS

### Animal Husbandry and Care

All animals were handled in accordance with the Association for Research in Vision and Ophthalmology (ARVO) statement for the use of animals in ophthalmic and vision research and monitored by the Institutional Animal Care and Use Committee (IACUC) of Columbia University.

Animals used in this study were 2-month-old males; C57/B6J females are protected from retinal edema. The tamoxifen inducible EC Casp9 mouse line was generated by crossing Cdh5(Pac)-Cre/ERT2 (Pitulescu, Schmidt, Benedito, & Adams, 2010) with caspase-9 flox/flox mice (Avrutsky et al., 2020). Tamoxifen was dissolved in corn oil (20 mg/mL), then 2mg of tamoxifen was administered intraperitoneally (IP) for five consecutive days when animals were eight weeks old (**Figure 1a**).

**Figure 1.**
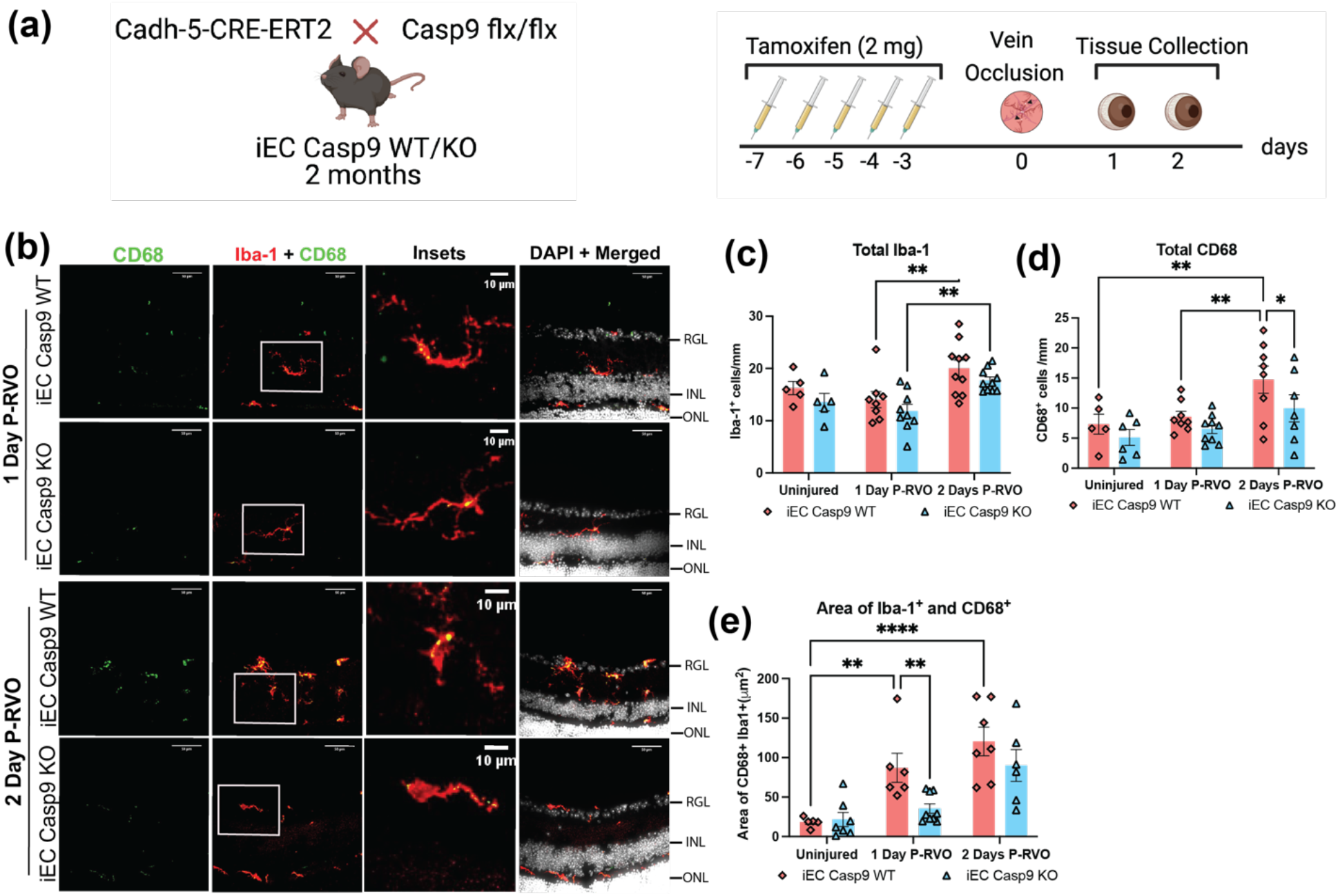
RVO induces an EC Casp9 regulated increase in CD68^+^ microglia. **(a)** Experimental schematic. Two-month-old iEC Casp9 WT/KO mice were treated with tamoxifen for five consecutive days. After two days, eyes were subjected to RVO, and eyes were collected one and two days P-RVO. Created with BioRender.com. **(b)** Retinal cross-sections from one and two days P-RVO iEC Casp9 WT and KO mice were stained with CD68 (green), Iba-1 (red), and DAPI (white). Scale bar 50μm and inset 10μm. **(c)** Quantification of the number of Iba-1^+^ cells in uninjured iEC Casp9 WT (n= 5) and KO (n= 6), one day P-RVO iEC Casp9 WT (n= 8) and KO (n= 9), and two days P-RVO iEC Casp9 WT (n= 10) and KO (n= 10). **(d)** Quantification of number of CD68^+^ cells in uninjured iEC Casp9 WT (n= 5) and KO (n= 6), one day P-RVO iEC Casp9 WT (n= 8) and KO (n= 9), and two days P-RVO iEC Casp9 WT (n= 8) and KO (n= 7). **(e)** Quantification of percent area of colocalization of Iba-1^+^ and CD68^+^ cells in uninjured iEC Casp9 WT (n= 5) and KO (n= 5), one day P-RVO iEC Casp9 WT (n= 6) and KO (n= 9), and two days P-RVO iEC Casp9 WT (n= 7) and KO (n= 6). Error bars mean ± SEM; Two-way ANOVA, Fisher’s LSD test. Retinal ganglion layer (RGL), inner nuclear layer (INL), and outer nuclear layer (ONL).

### Mouse Model of Retinal Vein Occlusion (RVO)

RVO was induced in major retinal veins (n=2-3 veins occluded/eye) 10 minutes after tail vein injection of the photoactivatable dye, Rose Bengal (37.5 mg/kg). Eight minutes after injection, eyes were dilated with tropicamide, and phenylephrine chloride eye drops, and mice were anesthetized with IP administration of a cocktail of ketamine (80-100 mg/kg) and xylazine (5-10 mg/kg). Two minutes after injection, depth of anesthesia was confirmed by toe-pinch. Afterwards, retinal veins were irradiated with the Micron IV image guided laser (532 nm) from Phoenix Research labs by delivering three adjacent laser pulses (power 100 mW, spot size spot size 50 μM, duration 1 second, total energy 0.3 J) to each vein at a distance of 375 μM from the optic nerve head. RVO occlusions were examined by fundus imaging after irradiation and one day after RVO (P-RVO). Occluded veins were identified by a whitening of the vessel at the site of the occlusion, dilation distal to the occlusion, and constriction of vessel diameter proximal to the occlusion. Persistent occlusions from the time of irradiation up to one day P-RVO, were classified as successful occlusions. Retinas with detachment, subretinal hemorrhage, or full ischemia one day P-RVO were excluded from further analysis. A detailed description of the RVO mouse model can be found in (Colon Ortiz, Potenski, Lawson, Smart, & Troy, 2021).

### Image Guided Optical Coherence Tomography (OCT) and Analysis

Following anesthesia and pupil dilation as indicated above in the mouse model of RVO section, OCT images were captured using the Phoenix Micron IV image-guided OCT system. Two vertical and two horizontal OCT scans were taken 75 μm from the periphery of the RVO burn areas, pre-RVO, and one day P-RVO. Total retinal thickness was determined using the InSight software.

### Disorganization of Inner Retinal Layers (DRIL)

Four OCT images per retina were analyzed in Image J for presence of DRIL. DRIL was measured with a horizontal line of indistinctive boundaries between the inner nuclear layer (INL) and the outer plexiform layer (OPL).

### Optokinetic Response Test

*In vivo* examination of the mouse optokinetic response was performed using the Striatech Optodrum. The awake mouse was placed on the platform inside the arena where the Optodrum presented a rotating stripe pattern to the animals at a rotation speed of 12°/second. Contrast sensitivity (1/contrast threshold) was determined by assessing the contrast threshold after presenting incremental visual acuity patterns at 0.05, 0.15 and 0.25 cycles/°. Readings were taken one day pre-RVO, one and two days P-RVO.

### Immunohistochemistry (IHC) and Analysis

Mice were perfused with sterile saline and then with 4% paraformaldehyde (PFA). Eyes were enucleated and immersed in 4% PFA overnight at 4°C, they were then washed three times for 10 minutes with 1X PBS and immersed in 30% sucrose for three nights at 4°C. Eyes were embedded in optimal cutting temperature solution and kept at −80°C. Embedded eyes were sectioned at 20 μm thickness. Retinal slides were chosen to match approximate location of OCT scans and then were washed with 1X PBS for 5 minutes. Sections were permeabilized with 0.1% Triton X-100 for two hours and blocked with blocking buffer (10% Normal Goat Serum (NGS), 1%Bovine Serum Albumin (BSA) dissolved in 1X PBS and filtered) overnight at 4°C. Primary antibodies for Iba-1 (1:200, BioCare CP29013), CD68 (1:100 BioRad MCA1957GA), cl-caspase-6 (1:100, Cell Signaling 9761S), GFAP (1:2,000 AVES GFAP), nestin (1:100, AVES Nes), and AQP-4 (1:100 Cell Signaling 59678S) were prepared with blocking buffer and sections were incubated overnight at 4°C. Afterwards, sections were washed four times with 1X PBS for five minutes, secondary antibodies were diluted in blocking buffer (1:1,000) and sections incubated for two hours at room temperature. Then, sections were washed four times with 1X PBS for five minutes and then stained with nuclear staining (Hoechst, 1:5,000 dilution) for five minutes. After one wash with 1X PBS for five minutes, retinal sections were mounted with Fluoromount-G. Retinas were imaged using a spinning disk confocal microscope (BioVision Technologies) and analyzed using FIJI image analysis software. All images quantified assessed 3-6 field of views in four retinal sections per eye that were averaged. The total cell number of microglia was determined based on morphology and presence of microglial marker. Microglial morphology was analyzed using the particle analysis function. Colocalization analysis was assessed with the color threshold function. Percent area of expression was determined by using the thresholding function.

### Western Blot and Analysis

Animals were perfused with saline and retinas were then dissected, minced, and placed in RIPA buffer with protease (Fisher) and phosphatase inhibitors II and III (Sigma). Tissue was then sonicated with a Sonic Dismembrator (Model 500, Fisher Scientific) for one minute (1 second on/ 1 second off). Retinas were matched one day P-RVO based on the fraction of veins occluded and disorganization of inner retinal layers (DRIL) percentages, two matched retinas were then combined. Protein concentrations were determined using a BCA assay. A total of 100μg of retinal protein was loaded per sample. Total protein transfer was determined using REVERT (LI-COR). Protein blots were blocked with LI-COR Blocking Buffer for one hour at room temperature and then incubated for two nights with primary antibody (GFAP GA5 Sigma 63893) at 4°C. Blots were washed three times with 0.1% TBS-Tween 20 and then incubated with secondary antibodies at 1:5,000 overnight at 4°C. After three ten-minute washes with 0.1% TBS-Tween 20, blots were imaged using the LI-COR Odyssey Imaging System. Blots were analyzed using Image Studio Lite Software; all signals were first normalized to total protein loaded and then to uninjured littermate control values.

### Cytokine Arrays and Analysis

Retinal tissue was prepared the same way as for Western Blot, except for the replacement of RIPA buffer with the manufacturer’s cell lysis buffer. Cytokine arrays were performed according to manufacturer’s protocol (Abcam, ab211069) and blots were imaged using the LI-COR Odyssey Imaging System and analyzed with Image Studio Lite Software.

### Procedures and Statical Analysis

The RVO model, *in vivo* and *in vitro* measures and analysis were done by investigators blinded to animal genotype. Eyes that presented with retinal detachment, cataract, excessive edema or intraretinal hemorrhage were excluded from all analyses and only eyes that had one or more occlusion by one day P-RVO were considered viable. GraphPad Prism and Excel were used for statistical analysis. One-way and Two-way ANOVA were used to determined statistical differences followed by Fisher’s LSD test; data is presented as mean ± SEM.

## RESULTS

### EC Casp9 modulates microglial CD68 in a time-dependent manner P-RVO

Microglial proliferation and reactivity are observed in retinal injury as an immune response that can have neuroprotective and/or neurodegenerative consequences (Silverman & Wong, 2018). In RVO, the number of microglial cells and levels of CD68 increases, and these responses have been linked to RGC death (Ebneter et al., 2017; Jovanovic et al., 2020). Thus, we first sought to investigate whether EC Casp9 regulates a microglial immune response in RVO pathology using iEC Casp9 KO and WT littermates (**Figure 1a**). To determine this, we assessed the total number of microglia by immunohistochemistry (IHC) with the microglial marker, Iba-1, of uninjured, one and two days P-RVO retinal sections (**Figure 1b**). Quantification analysis showed that the total number of Iba-1^+^ cells increases significantly two days P-RVO when comparing one day P-RVO to two-days P-RVO retinas, independent of EC Casp9 expression (**Figure 1c**). We then evaluated the expression of the microglial lysosomal marker, CD68, that is associated with a microglial inflammatory state and is increased in RVO (Ebneter et al., 2017; Hirabayashi et al., 2019), by quantifying the total number of CD68^+^ cells. The data indicate that RVO induces a significant increase in the number of CD68^+^ cells by two days P-RVO compared to uninjured and one day P-RVO iEC Casp9 WT retinas. Knockout of EC Casp9 led to a significant decrease in total number of CD68^+^ cells compared to injured iEC Casp9 WT retinas two days P-RVO (**Figure 1d**). To further determine the level of CD68 in microglial cells, we analyzed the area of colocalization between Iba-1 and CD68. The results demonstrate that RVO induces a significant upregulation of CD68 in microglia one and two days P-RVO, compared to uninjured iEC Casp9 WT retinas. EC Casp9 deletion led to significantly less induction of CD68 in Iba-1 cells one day P-RVO, a trend that is sustained through two days P-RVO (**Figure 1e**). Furthermore, we assessed whether RVO modulated microglial morphology in the context of EC Casp9. Our analysis indicates that microglia become bigger, hypertrophic and more circular one day P-RVO compared to uninjured iEC Casp9 WT retinas, and these changes in area and Ferret’s diameter are blocked by deletion of EC Casp9 one day P-RVO, but by two days P-RVO there is no significant difference between genotypes. At two days P-RVO the change in microglia circularity decreases in both genotypes (**Figure S1**).

### EC Casp9 leads to macroglial changes in nestin and AQP-4 P-RVO in a time-dependent manner

Other markers of macrogliosis include nestin and aquaporin-4 (AQP-4). Nestin is an intermediate filament whose expression is downregulated in mature macroglia (Gilyarov, 2008). In hypoxic-ischemic injury, nestin increases and leads to changes in macroglial motility and cell division (Cho, Shin, Park, Kim, & Lee, 2013; Lin, Matesic, Marvin, McKay, & Brustle, 1995). AQP-4 regulates a water channel protein which levels are implicated in retinal vascular disease (Cui, Sun, Xiang, Liu, & Li, 2012; Nicchia et al., 2016). To determine if RVO and EC Casp9 modulated components that are known markers for macrogliosis, we assessed the levels of nestin and AQP-4 by IHC analysis and quantification in the RGL and NFL, and in the whole retina by evaluating the percent area of expression. Data analysis shows that RVO induces a significant increase in the level of nestin in the RGL and NFL, and whole retina two days P-RVO (**Figure 2a,b**), and that EC Casp9 deletion abrogates the increase in whole retina, but does not alter the increase in RGL and NFL (**Figure 2a,c**). Analysis of AQP-4 levels in the RGL and NFL and whole retina,demonstrates that RVO induces a significant decrease in its level by one day P-RVO and that its levels are restored back to uninjured levels by two days P-RVO in the injured iEC Casp9 WT retinas. Levels of AQP-4 in the iEC Casp9 KO were significantly higher than in the iEC Casp9 WT two days P-RVO (**Figure 2d-f**).

**Figure 2.**
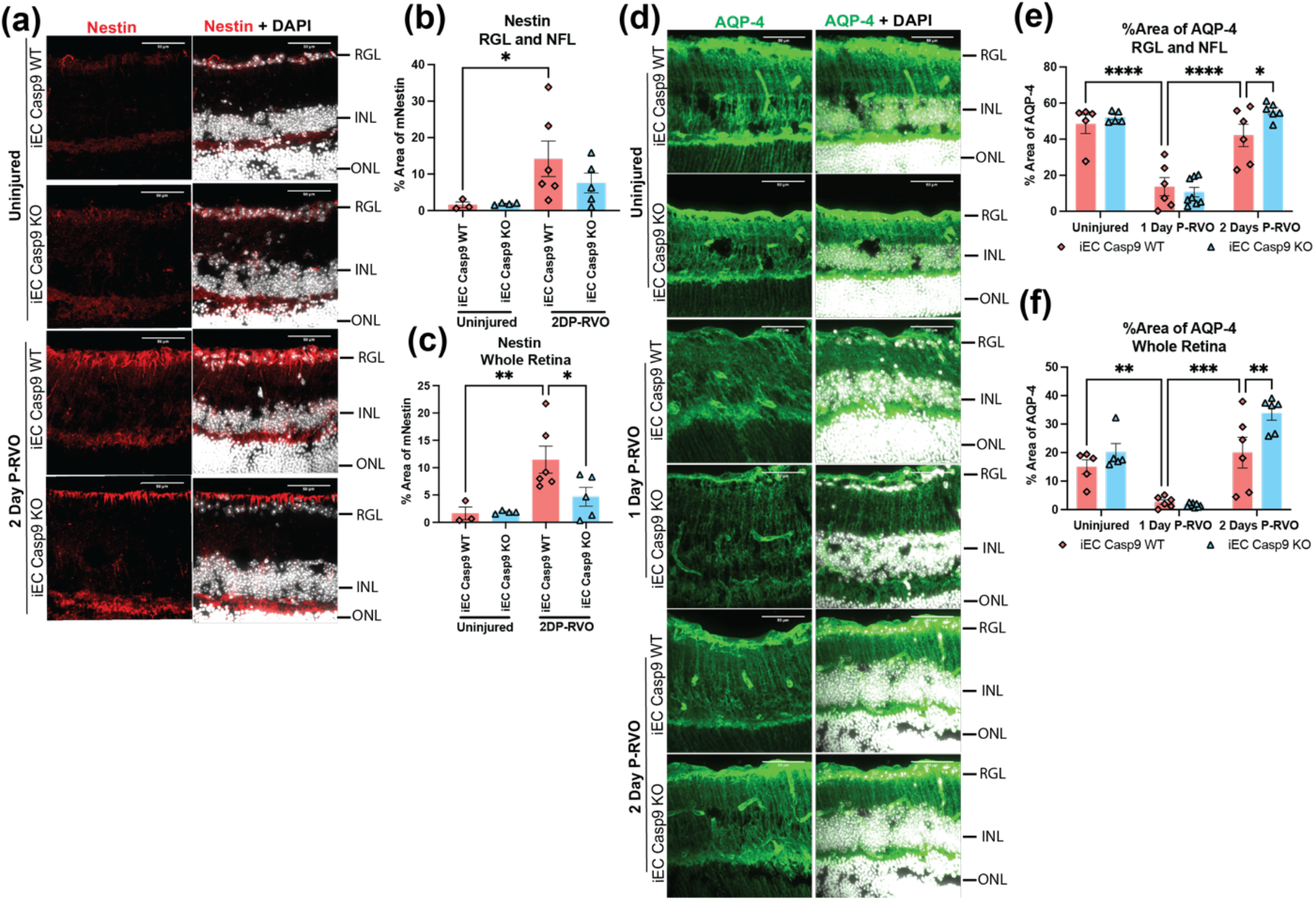
EC Casp9 induces macroglial changes in nestin and AQP-4 two days P-RVO. **(a)** Retinal cross-sections from uninjured and two days P-RVO iEC Casp9 WT and KO mice were stained with nestin (red) and DAPI (white). Scale bar=50μm. **(b)** Quantification of percent area of expression of nestin in RGL and NFL of uninjured iEC Casp9 WT (n= 3) and KO (n= 5) and two days P-RVO iEC Casp9 WT (n= 6) and KO (n= 5). **(c)** Quantification of percent area of expression of nestin in whole retina of uninjured iEC Casp9 WT (n= 3) and KO (n= 5) and two days P-RVO iEC Casp9 WT (n= 6) and KO (n= 5). Error bars mean ± SEM; One-way ANOVA, Fisher’s LSD test. **(d)** Retinal cross-sections of uninjured, one and two-days P-RVO iEC Casp9 WT and KO stained with AQP-4 (green) and DAPI (white). Scale bar=50μm. **(e)** Quantification of percent area of expression of AQP-4 in RGL and NFL of uninjured iEC Casp9 WT (n= 5) and KO (n= 5) one day P-RVO iEC Casp9 WT (n= 6) and KO (n= 8), and two days P-RVO iEC Casp9 WT (n= 6) and KO (n= 6). **(f)** Quantification of percent area of expression of AQP-4 in whole retina of uninjured iEC Casp9 WT (n= 6) and KO (n= 8) one day P-RVO iEC Casp9 WT (n= 10) and KO (n= 6), and two days P-RVO iEC Casp9 WT (n= 6) and KO (n= 8). Error bars mean ±SEM; Two-way ANOVA, Fisher’s LSD test. Retinal ganglion layer (RGL), inner nuclear layer (INL), and outer nuclear layer (ONL).

### EC Casp9 increases astroglial cl-caspase-6 and contributes to caspase-6 induced GFAP cleavage P-RVO

Prior work showed that pharmacologically inhibiting caspase-9 decreased the expression of its downstream target caspase-6 in astrocytes P-RVO (Avrutsky et al., 2020). To evaluate if EC Casp9 specifically modulated cl-caspase-6, we stained uninjured, one and two days P-RVO sectioned retinas from iEC Casp9 KO and WT littermates and then evaluated the level of cl-caspase-6 by quantifying the percent area of expression in the RGL and NFL, areas where astrocytes are located in the retina. The analysis revealed that RVO induces an increase in cl-caspase-6 two days after injury and that EC Casp9 deletion significantly blocks the change in cl-caspase-6 levels in the RGL and NFL one and two days P-RVO (**Figure 3a-b**). Similar trends were also noted in the analysis of the level of cl-caspase-6 in the inner nuclear layer (INL), indicating an increase of cl-caspase-6 in the retinal neurons, which is blocked by EC Casp9 deletion two days P-RVO (**Figure 3c**).

**Figure 3.**
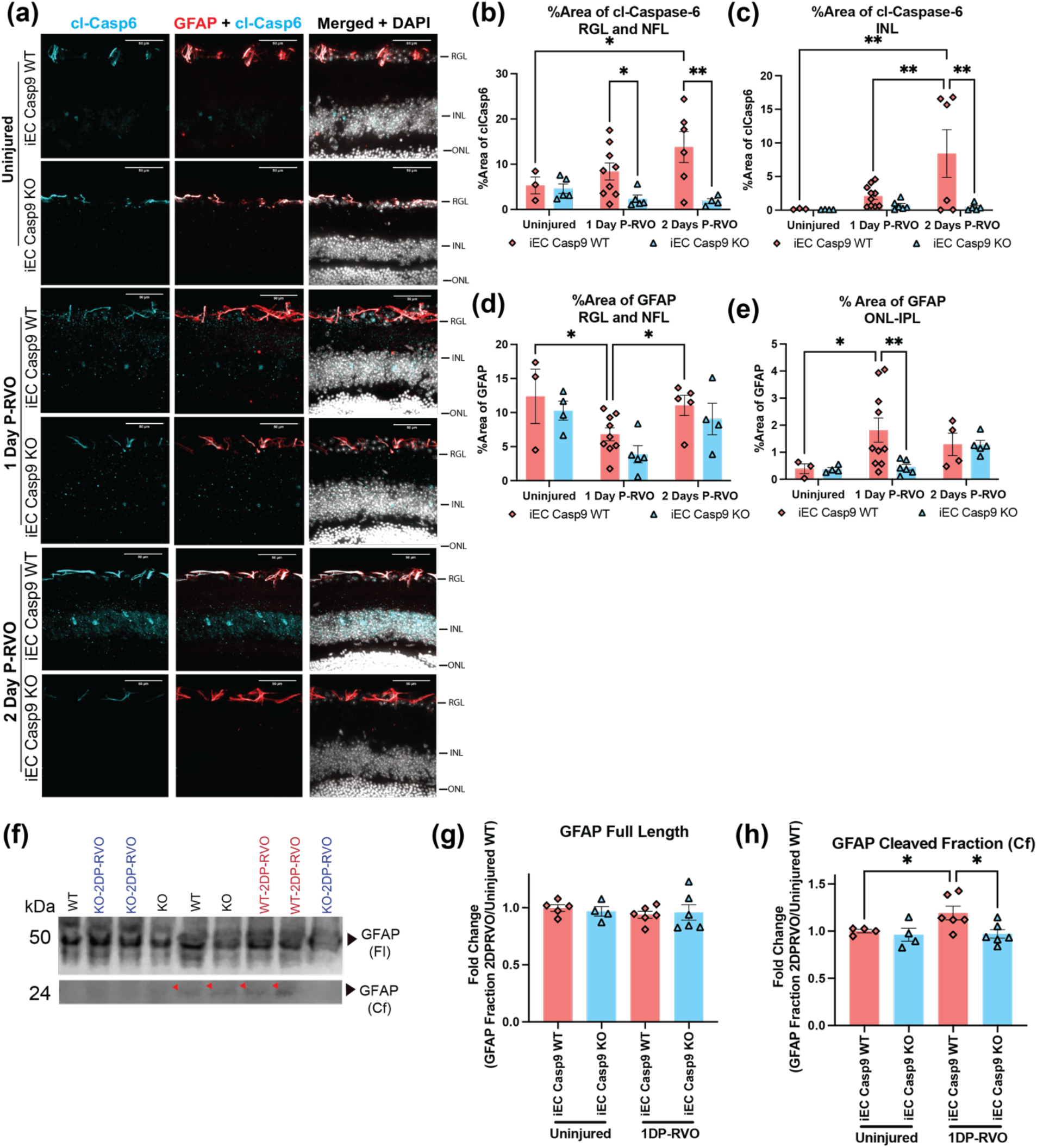
EC Casp9 promotes astroglial increase in cl-caspase-6 levels and caspase-6 GFAP cleavage. **(a)** Retinal cross-sections from one and two days P-RVO iEC Casp9 WT and KO mice were stained with cl-caspase-6 (cyan), GFAP (red), co-localization of cl-Casp6 and GFAP (white), and DAPI (white). Scale bar=50μm. **(b)** Percent area of expression of cl-caspase-6 in RGL and NFL of uninjured iEC Casp9 WT (n= 3) and KO (n= 5), one day P-RVO iEC Casp9 WT (n= 9) and KO (n= 6), and two days P-RVO iEC Casp9 WT (n= 6) and KO (n= 4). **(c)** Percent area of expression of cl-caspase-6 in INL of uninjured iEC Casp9 WT (n= 3) and KO (n= 5), one day P-RVO iEC Casp9 WT (n= 10) and KO (n= 6), and two days P-RVO iEC Casp9 WT (n= 6) and KO (n= 5). **(d)** Percent area of expression of GFAP in RGL and NFL of uninjured iEC Casp9 WT (n= 3) and KO (n= 4), one day P-RVO iEC Casp9 WT (n= 9) and KO (n= 5), and two days P-RVO iEC Casp9 WT (n= 4) and KO (n= 6). **(e)** Percent area of expression of GFAP from ONL to IPL in iEC Casp9 WT (n= 3) and KO (n= 4), one day P-RVO iEC Casp9 WT (n= 10) and KO (n= 6), and two days P-RVO iEC Casp9 WT (n= 4) and KO (n= 5). Error bars mean ± SEM; Two-way ANOVA, Fisher’s LSD test. Retinal ganglion layer (RGL), inner nuclear layer (INL), and outer nuclear layer (ONL). **(f)** Western blot detection of GFAP in uninjured iEC Casp9 WT and KO and two days P-RVO iEC Casp9 WT and KO retinal lysates. **(g)** Quantification of GFAP full length 50 kDa band in uninjured iEC Casp9 WT (n=5, biological replicates) and KO (n=4, biological replicates) and two-days P-RVO WT (n=6, biological replicates) and KO (n=6, biological replicates) normalized to total protein and uninjured iEC Casp9 WT. **(h)** Quantification of GFAP cleaved fraction 24 kDa in uninjured iEC Casp9 WT (n=5, biological replicates) and KO (n=4, biological replicates) and two-days P-RVO WT and KO normalized to total protein and uninjured iEC Casp9 WT. Error bars mean ± SEM; One-way ANOVA, Fisher’s LSD test.

Next, we sought to determine if RVO and EC Casp9 induced changes in GFAP, as increased levels of GFAP are often associated with an inflammatory response of Müller glia and astrocytes (Eng & Ghirnikar, 1994). IHC analysis of GFAP in the RGL and NFL suggested that there are no changes in the level of GFAP at early time points in RVO nor are GFAP levels modified by deletion of EC Casp9 (**Figure 3d**). However, analysis of GFAP levels from the outer nuclear layer (ONL) to the inner plexiform layer (IPL), showed a significant increase of GFAP in injured iEC Casp9 WT one day P-RVO that is modulated by deletion of EC Casp9. No differences were noted in the expression of GFAP from ONL to IPL by two days P-RVO (**Figure 3e, Figure S2**).

In injury models GFAP can be cleaved by caspase-6 (Battaglia et al., 2019; Jonesco et al., 2019; Mouser, Head, Ha, & Rohn, 2006). To test whether GFAP is a substrate of caspase-6 in RVO, we performed a western blot analysis two days P-RVO. For each sample, two retinas were matched by similar fractions of veins occluded and percentage of disorganization of inner retinal layers (DRIL), an indicator of capillary ischemia. Quantification of the GFAP 50 kDa band demonstrated that there were no differences in injured samples compared to uninjured regardless of genotype (**Figure 3f,g**). However, analysis of the GFAP caspase-6 cleaved fraction, the 24 kDa band, showed that RVO induced a significant increase in GFAP cleavage in EC Casp9 WT compared to uninjured. This increase in GFAP cleavage was reduced to baseline levels in injured EC Casp9 KO (**Figure 3f,h**).

### EC Casp9 contributes to changes in the levels of inflammatory cytokines one day P-RVO

Increased levels of inflammatory cytokines in RVO patients correlate with hypoxic ischemic injury and retinal edema (Noma et al., 2014). Our previous work indicated that the peak of retinal edema for the EC Casp9 mouse line is two days P-RVO (Avrutsky et al., 2020). We collected retinas one day P-RVO to determine which inflammatory cytokines could precede the peak of edema by performing a cytokine array using two retinas per sample as described above. We found that RVO led to a significant increase of the following cytokines in the WT retinas: CX3CL1 (fractalkine), insulin growth factor-1 (IGF-1), IL-4, LIX (CXCL5), IL-1α, macrophage colony stimulating factor (M-CSF), tumor necrosis factor (TNF-α), IL-1β, IL-10, and vascular endothelial growth factor A (VEGF-A); the cytokine changes were attenuated in injured EC Casp9 KO retinas (**Figure 4a-l**). Other cytokines that we found to be upregulated in RVO independently of EC Casp9 expression were MMP3 and MCP1 (**Figure 4m-n**). We also found that MMP-2, SDF-1α, and TGF-β were decreased in injured iEC Casp9 KO compared to WT but these cytokines were not significantly modulated by RVO in Casp9 WT animals (**Figure S3**).

**Figure 4.**
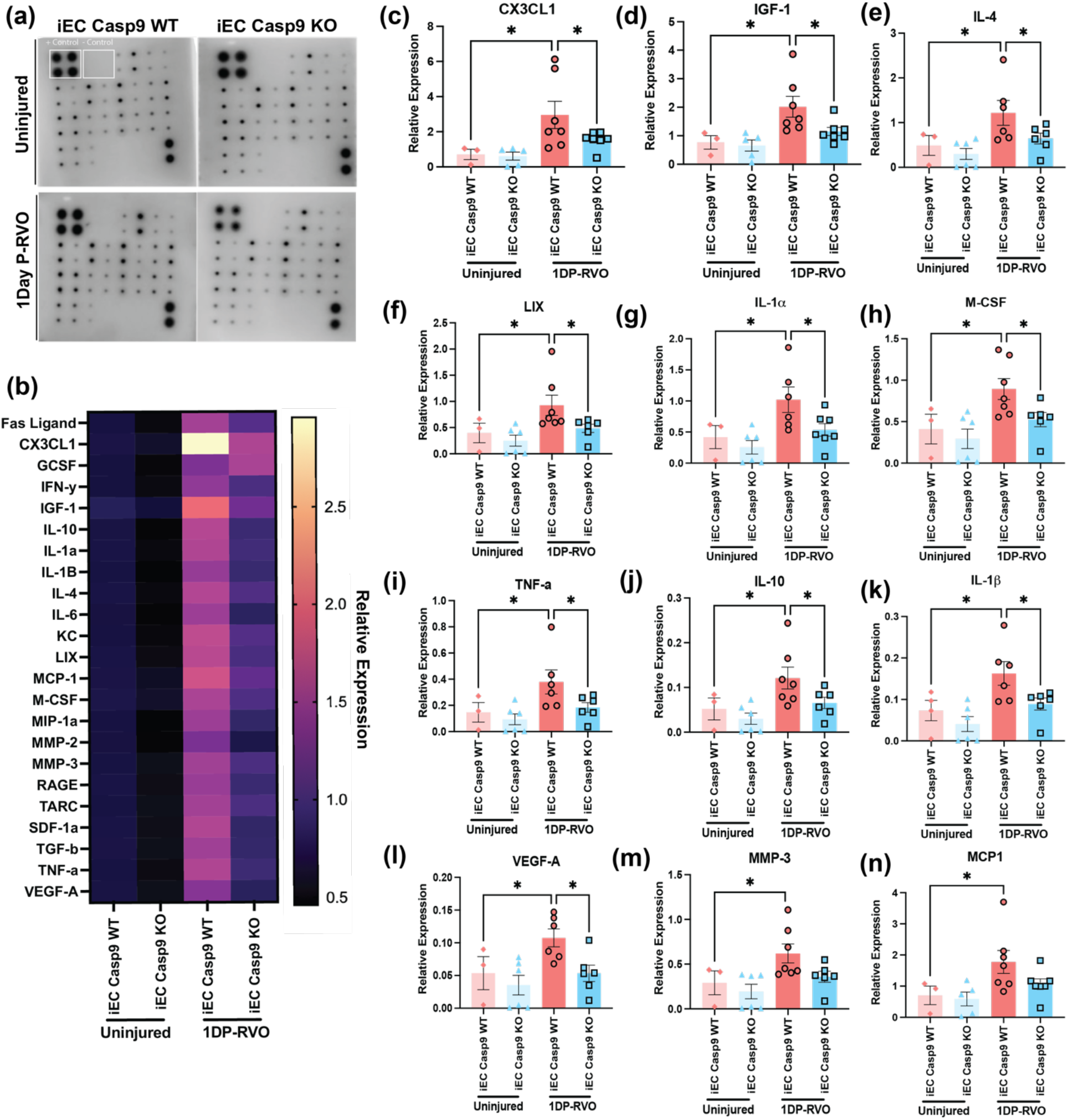
EC Casp9 loss decreases expression of cytokines one day P-RVO. **(a)** Representative images of cytokine arrays of retinas from uninjured iEC Casp9 WT/KO and one day P-RVO iEC Casp9 WT/KO mice. **(b)** Heat Map array of relative expression of 23 cytokines normalized to uninjured iEC Casp9 WT in uninjured iEC Casp9 WT (n= 3) and KO (n= 6) and iEC Casp9 WT (n= 6) and KO (n= 6) one day P-RVO. **(c-n)** Relative expression of induced cytokines one day P-RVO. Error bars mean ± SEM; One-way ANOVA, Fisher’s LSD test.

We then assessed potential interactions of caspase-9 with RVO induced and EC Casp9 regulated cytokines by performing STRING (Szklarczyk et al., 2019) and gene ontology (GO) analysis. The analysis suggests potentially significant interactions in the network, pinpointing experimentally determined connections of caspase-9 with the TNF-superfamily and IL-1β (**Figure 5a**, pink lines). Other potential caspase-9 interactions are suggested with IL-10, IGF-1, and VEGF-A through textmining (**Figure 5a**, green lines). Some of the most significant biological processes revealed were related to neuronal death, regulation of immune system processes, response to external stimulus, and leukocyte migration (**Figure 5b**). When looking at the strongest biological associated processes, some are related to chronic inflammatory response to antigenic stimulus, and positive regulation of microglial cell migration (**Figure 5c**). Similar levels of significant interactions were found when we evaluated the caspase-9 downstream targets, caspase-7 and caspase-6 using STRING analysis (**Figure S4**).

**Figure 5.**
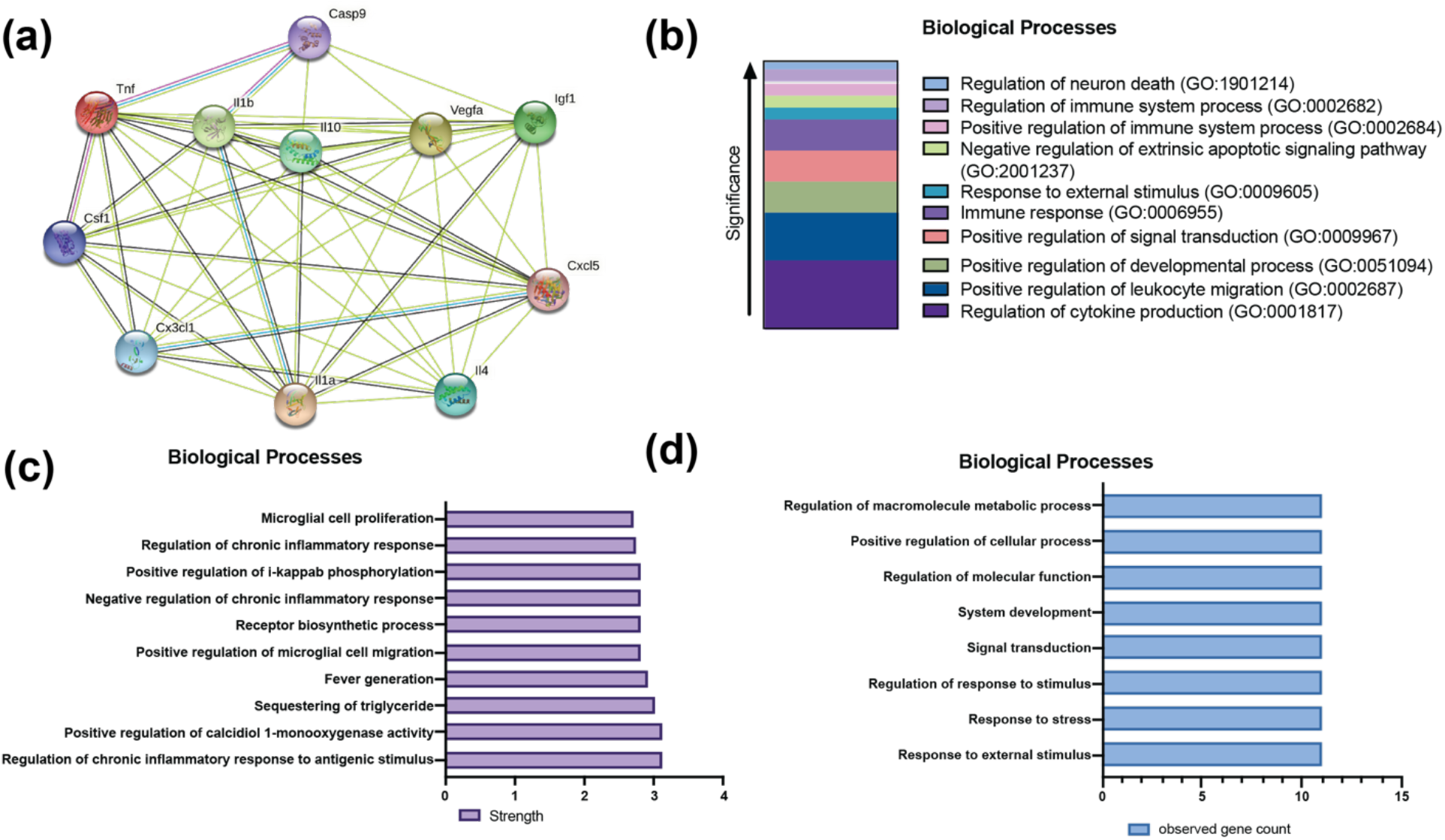
STRING analysis of cytokines modified by EC Casp9. **(a)** Protein-protein network interaction of cytokines regulated by EC Casp9 P-RVO. **(b)** Gene ontology analysis of biological processes associated with cytokines based on significance of the enrichment. **(c)** Strength of the gene ontology analysis of biological processes. **(d)** Biological terms associated with all cytokines modulated by EC Casp9.

### EC Casp9 promotes contrast sensitivity decline P-RVO

One of the effects of RVO is damage to visual function, including a significant decline in contrast sensitivity threshold (McIntosh et al., 2010; Mishra et al., 2020). Since we had previously found that EC Casp9 regulated retinal edema and retinal neuronal death in RVO (Avrutsky et al., 2020), we sought to explore whether EC Casp9 was implicated in damage to contrast sensitivity discrimination. To this end, we evaluated contrast sensitivity with incremental visual acuity ranges of 0.05, 0.15, and 0.25 cycles/° (**Figure 6a-b**). But first, we probed if there were any differences in visual function between uninjured iEC Casp9 WT and iEC Casp9 KO mice by performing separate contrast sensitivity and visual acuity tests (**Figure 6c-d**). The results suggest that there are not any differences in visual function performance between genotypes and we then proceeded with to test contrast sensitivity discrimination with the incremental visual acuity ranges. We found that RVO causes a significant decrease in contrast sensitivity at 0.05 cycles/° and this loss was rescued one day P-RVO in animals that lacked EC Casp9 (**Figure 6e**). The degree of contrast sensitivity significantly correlated with the fractions of veins occluded in iEC Casp9 WT but not in iEC Casp9 KO animals **(Figure 6f**).

**Figure 6.**
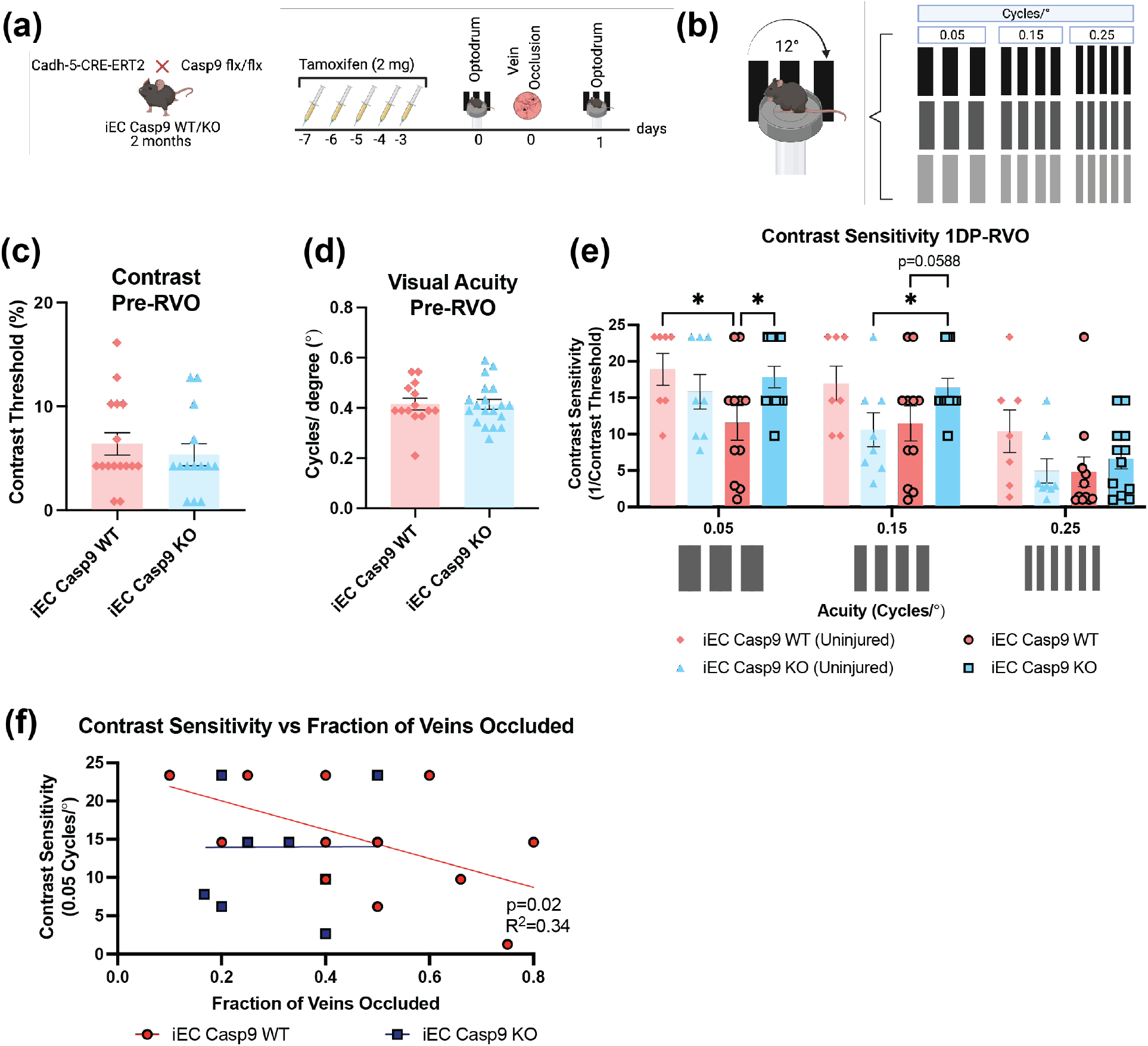
EC Casp9 loss rescues contrast sensitivity one day P-RVO. **(a)** Experimental schematic. Two-month-old iEC Casp9 WT/KO mice were treated with tamoxifen for five consecutive days. After two days of rest, their optokinetic response was tested pre and one day P-RVO. Created with BioRender.com. **(b)** Optokinetic test was performed by testing their contrast sensitivity at three acuity settings: 0.05, 0.15, 0.25 cycles/°. **(c-d)** Contrast sensitivity and visual acuity values prior to RVO of uninjured iEC Casp9 WT (n=14-15) and KO (n=14-19). **(e)** Contrast sensitivity response of uninjured iEC Casp9 WT (n=7) and KO (n=8), and iEC Casp9 WT (n= 11) and KO (n= 12) one day P-RVO at 0.05, 0.15, 0.25 cycles/°. Error bars mean ± SEM; Two-way ANOVA, Fisher’s LSD test. **(f)** Simple linear regression of contrast sensitivity at 0.05 cycles/° vs fraction of veins occluded at one day P-RVO.

## DISCUSSION

We have previously shown that non-apoptotic EC Casp9 signaling regulates many of the retinal changes linked to retinal vascular injury, and now queried whether this signaling was also required for impairment of vison and inflammation. In this study, we show that caspase-9 in hypoxic endothelial cells regulates changes in visual function and retinal inflammatory signaling in microglia, astrocytes and Müller cells. By using an established mouse model of RVO, we found that EC Casp9 causes an increase in: (1) reactive microglia, (2) inflammatory cytokines, and (3) cleaved-caspase-6 and GFAP cleavage in astrocytes. EC Casp9 also regulates (4) changes in GFAP, nestin and AQP4 levels in Müller glia, and (5) contrast sensitivity decline (**Figure 7**).

**Figure 7.**
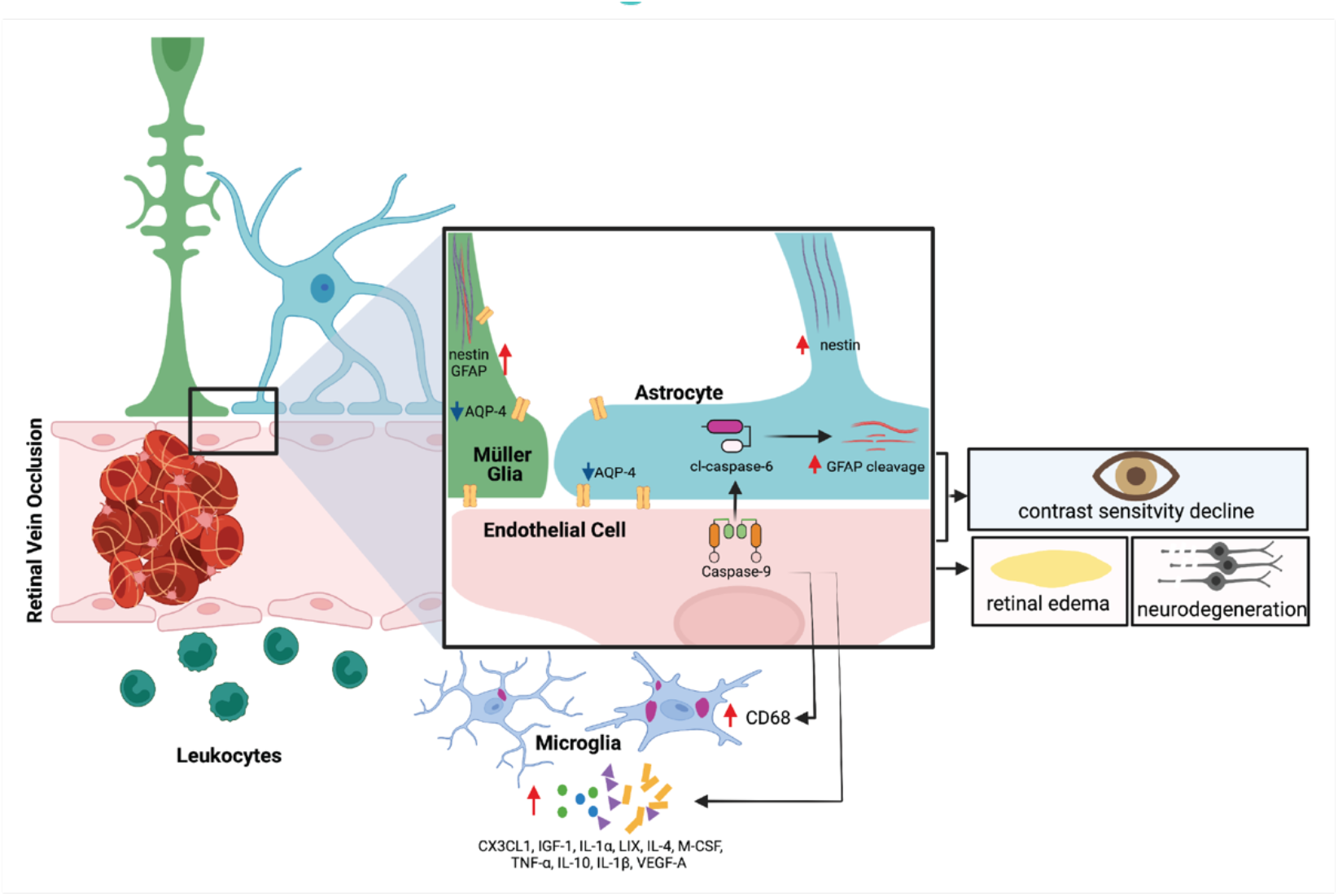
Model of EC Casp9 regulation of glia and cytokines in RVO. Created with BioRender.com.

In a model of branch RVO (occlusion of one retinal vein) microglia respond by clustering and increasing in number near the site of injury three days P-RVO, with a peak in microglial number at seven days (Ebneter et al., 2017). Our study revealed that RVO leads to an increase in microglial number as early as two-days P-RVO independent of EC Casp9. To better visualize and appreciate individual microglial cells and their localization in respect to retinal layers, we used retinal sections for microglial analysis. One drawback of this technique is that it does not allow for analysis of microglial clusters and mobilization towards the occluded veins. The identification of pro-inflammatory microglia has been ascribed to morphological changes from ramified to ameboid in the context of disease or injury (Giulian, 1987). We found that RVO changes the morphology of microglia towards a bigger, more circular, and hypertrophic shape. This is the first study to quantify RVO-induced microglial morphological changes. However, EC Casp9 regulation of microglial morphology was transient as it was only present at one day P-RVO. It is important to note that a limitation of the microglial morphological analysis used in this study is that it did not target all microglial cells or processes, but this was consistent in all the images that were evaluated.

It is known that microglial shape is linked to function, but changes in shape are not sufficient to understand their potential pro- or anti-inflammatory pathological function which can be context dependent (Davis, Salinas-Navarro, Cordeiro, Moons, & De Groef, 2017; Fernandez-Arjona, Grondona, Fernandez-Llebrez, & Lopez-Avalos, 2019). Therefore, markers such as CD68 and F4/80 are commonly used to histologically survey the population of reactive or pro-inflammatory microglial cells. CD68 is a macrosialin receptor localized to the lysosome of macrophages/microglia and can be used to determine their phagocytic potential (Graeber, Streit, Kiefer, Schoen, & Kreutzberg, 1990). Like other reports (Ebneter et al., 2017; Hirabayashi et al., 2019), we found that RVO induces an increase in the number of CD68^+^ cells –which could be infiltrating monocytes in addition to microglia– and levels of microglial CD68. In RVO, this microglial phenotype potentially leads to phagocytosis of debris and clearance of apoptotic bodies to regulate continued inflammation and neuronal apoptosis (Marquez-Ropero, Benito, Plaza-Zabala, & Sierra, 2020). Our data shows that EC Casp9 deletion reduced levels of microglial CD68 at one day P-RVO and the number of CD68^+^ cells by two days P-RVO. The microglial changes mediated by EC Casp9 (microglial area, hypertrophy and levels of microglial CD68 at one day P-RVO and number of CD68^+^ cells at two days P-RVO) elucidates that there are distinctive microglia clusters that could mediate the development of retinal edema at early time points. A potential mechanism by which microglia contribute to retinal edema is through the release of inflammatory cytokines. Many of the cytokines we found to be modulated by the depletion of EC Casp9 are mostly expressed by microglial cells (Zhang et al., 2014; Zhang et al., 2016). It has been shown that depletion of microglia and leukocytes in RVO led to a significant decrease in inflammatory cytokines (Jovanovic et al., 2020). Evaluating the results of our study and Jovanovic et al., 2020, we can infer that in RVO some of the cytokines regulated by EC Casp9 are more likely to come from microglia. A limitation of our study is that we do not know which ccells are expressing the EC Casp9 modulated cytokines.

Pro-inflammatory cytokines are known to play an important role in RVO as they correlate with macular edema and hypoxic-ischemic injury (Noma et al., 2014). Here, we found that EC Casp9 signaling regulated several pro-inflammatory cytokines (IGF-1, IL-1α, IL-1β, M-CSF, TNF-α, and VEGF-A), anti-inflammatory cytokines (IL-4 and IL-10), and chemokines (CX3CL1 and LIX). A study that evaluated the levels of cytokines and chemokines in vitreous samples from ischemic RVO human patients showed increased levels of CX3CL1, TNF-α, IL-1β, IL-4, and IL-10 (Zeng et al., 2019), which we found to be modulated by EC Casp9. This evidence highlights the translational relevance of our model system. The importance of a more detailed understanding of the inflammatory pathways initiated by EC Casp9 points out key cytokines that could contribute to the development of retinal edema and neurodegeneration which could be targeted with therapeutic potential. While caspase-9 has not been shown to directly activate cytokines, downstream caspase-9 targets such as caspase-6, caspase-7 and caspase-3 are known to cleave cytokine proforms and increase overall cytokine levels (Harald Loppnow, 2000). Caspase-9 mediated activation of caspase-6 could contribute to the increased levels of TNF-α in RVO. When caspase-6 is activated, it can stimulate transcription and cause macrophage release of TNF-α (Ladha et al., 2018; Zheng, Karki, Vogel, & Kanneganti, 2020). Additionally, studies show that caspase-6 null mice have a significant decrease in TNF-α (Ladha et al., 2018). Many of the EC Casp9 regulated cytokines are target genes of the transcription factor NFκ-B (https://www.bu.edu/nf-kb/gene-resources/target-genes/). We found that EC Casp9 induced caspase-6 expression in neurons and astrocytes P-RVO, suggesting non-cell autonomous functions of EC Casp9. A recent *in vitro* enzymatic study suggested that NFκ-B is a potential substrate of caspase-6 (Julien et al., 2016). These data suggests that the EC Casp9 regulation of caspase-6 could lead to NFκ-B activation and subsequent mediation of pro-inflammatory cytokines in astrocytes – where we found the most expression of caspase-6 P-RVO.

Whether astrocytes become “reactive” P-RVO is a topic of controversy, with only a few studies assessing the levels of GFAP and AQP-4 (Koferl et al., 2014; McAllister et al., 2018; Rehak et al., 2009). Consistent with some of these studies (Koferl et al., 2014; Rehak et al., 2009), we found no differences in GFAP levels in the RGL and NFL at early timepoints P-RVO. When we evaluated the expression of another intermediate filament, nestin, we saw differences in the areas of the retina that were regulated by EC Casp9 P-RVO. Animals lacking EC Casp9 had less GFAP and nestin in the whole retina but not in the RGL and NFL. The difference in spatial expression of GFAP and nestin suggests that EC Casp9 potentially mediates Müller glia responses, as the RGL and NFL also contain the end feet of Müller glia. Therefore, the measured expression of GFAP and nestin in the RGL and NFL in this study could also indicate potential EC Casp9 induced changes in Müller glia that should be further assessed. We also found that RVO induces an increase in the level of cl-caspase-6 in astrocytes and cleaved GFAP that is abrogated by the deletion of EC Casp9. Caspase-6 cleavage of GFAP is known to cause GFAP hyper-aggregation and filament malformation (Chen, Hagemann, Quinlan, Messing, & Perng, 2013), and is associated with neurodegenerative diseases such as Alexander’s and Alzheimer’s disease (Battaglia et al., 2019; Mouser et al., 2006). However, the immunohistochemical techniques that we used were not able to detect GFAP hyper-aggregation or filament malformation histologically, but future studies could assess this possibility in RVO. Caspase-6 could be activated in astrocytes by caspase-9 or by self-dimerization, and how EC Casp9 leads to increased astroglial cl-caspase-6 is an area to explore. Potential endothelial-astroglia signaling pathways could involve neuronal release of apoptotic extracellular vesicles or endothelial release of molecules that can directly induce an astroglial response. This is the first study that identifies RVO-induced changes in caspase-6 expression in astrocytes.

The increase of proinflammatory cytokines in RVO correlates with a decrease in visual acuity (Jung, Kim, Sohn, & Yang, 2014; Kaneda et al., 2011). Thus, early changes in visual function P-RVO could be a result of the EC Casp9 regulated immune changes that could contribute to neuroretinal death. Patients with RVO present with poor visual acuity at the time of diagnosis, and in some cases vision can improve with time, but the damage significantly affects vision-related quality of life as described in the analysis of the visual function questionnaire (VFQ) reports (Awdeh et al., 2010; Deramo, Cox, Syed, Lee, & Fekrat, 2003). RVO also impacts contrast sensitivity, which has prospective use as indicator of disease progression (Mishra et al., 2020; Shoshani et al., 2011). Contrast sensitivity measurements allow the detection of changes in retinal blood flow (Shoshani et al., 2011) in retinal vascular disease, serving as a good tool for detecting early disease pathology and treatment efficacy. Contrast sensitivity is a reflection of RGC functionality as RGC are part of the contrast sensitivity pathway (Enroth-Cugell & Robson, 1966). We found that RVO-induced contrast sensitivity decline was rescued one day P-RVO when EC Casp9 was deleted. The EC Casp9 contribution to contrast sensitivity decline is consistent the role of EC Casp9 in mediating retinal edema and neuronal death (Avrutsky et al., 2020). Understanding the timing of vision contrast decline P-RVO and the role of EC Casp9 can help elucidate mechanisms of neurodegeneration. The observed contrast sensitivity decline could be due to degeneration and death of RGC types from the magnocellular or parvocellular vision pathways, as these types of RGC process contrast sensitivity (Silveira et al., 2004). RGC are vulnerable and degenerate in RVO (Alshareef et al., 2016), but the specific types of RGC that are most prone to degeneration in RVO have not been identified. Contrast sensitivity decline can also occur preceding degeneration and be indicative of RGC synaptic dysfunction.

We demonstrate an important role for EC Casp9 signaling in mediating inflammatory responses, which contributes to visual dysfunction in retinal neurovascular injury. These findings put into perspective the need to understand which neuroimmune responses are responsible for retinal edema and visual dysfunction, and could help develop therapeutics. Additionally, this study points out that detecting changes in contrast sensitivity at high spatial stimulus could be used as a biomarker to detect retinal vascular injuries.

## AUTHOR CONTRIBUTION

CCO performed the experiments, analyzed data, designed experiments, and wrote the manuscript. AMN performed experiments, analyzed data, and contributed to editing the manuscript. MIA performed the mouse model of RVO for one day P-RVO and helped with editing the manuscript. MC, JS, and JL performed surgery, optokinetic testing, animal husbandry and genotyping. CMT contributed to experimental design, funding acquisition, writing, and reviewing of the manuscript.

## ACKNOWLEDGEMENTS

We would like to thank Natasha Snider for kindly providing the GFAP GA5 antibody along with guidance and recommendations. We would also like to thank Marc Tessier-Lavigne and Ralf Adams for providing the caspase-9 flox/flox and the Cdh5(PAC)-CreERT2 mice, respectively. We would like to thank James Goldman for helpful discussions and suggestions. The research presented in this publication was supported by the National Science Foundation Graduate Research Fellowship Program (NSF-GRFP) grant DGE – 1644869, the National Institute of Neurological Disorders and Stroke (NINDS) of the National Institutes of Health (NIH), award number F99NS124180 NIH NINDS Diversity Specialized F99 (to CKCO), award number R03NS099920 (to CMT), and the Department of Defense Army/Air Force (DURIP to CMT). The content is the responsibility of the atuhors and does not represent the views of the National Institutes of Health.

## CONFLICT OF INTEREST STATEMENT

The authors declare the following conflict of interest: CMT and MIA are inventors on patent US 17/515,202. The remaining authors declare no conflict of interest.

## DATA SHARING STATEMENT

The data that support the findings of this study are available from the corresponding author upon reasonable request.

---

**Supplementary Figure 1.**
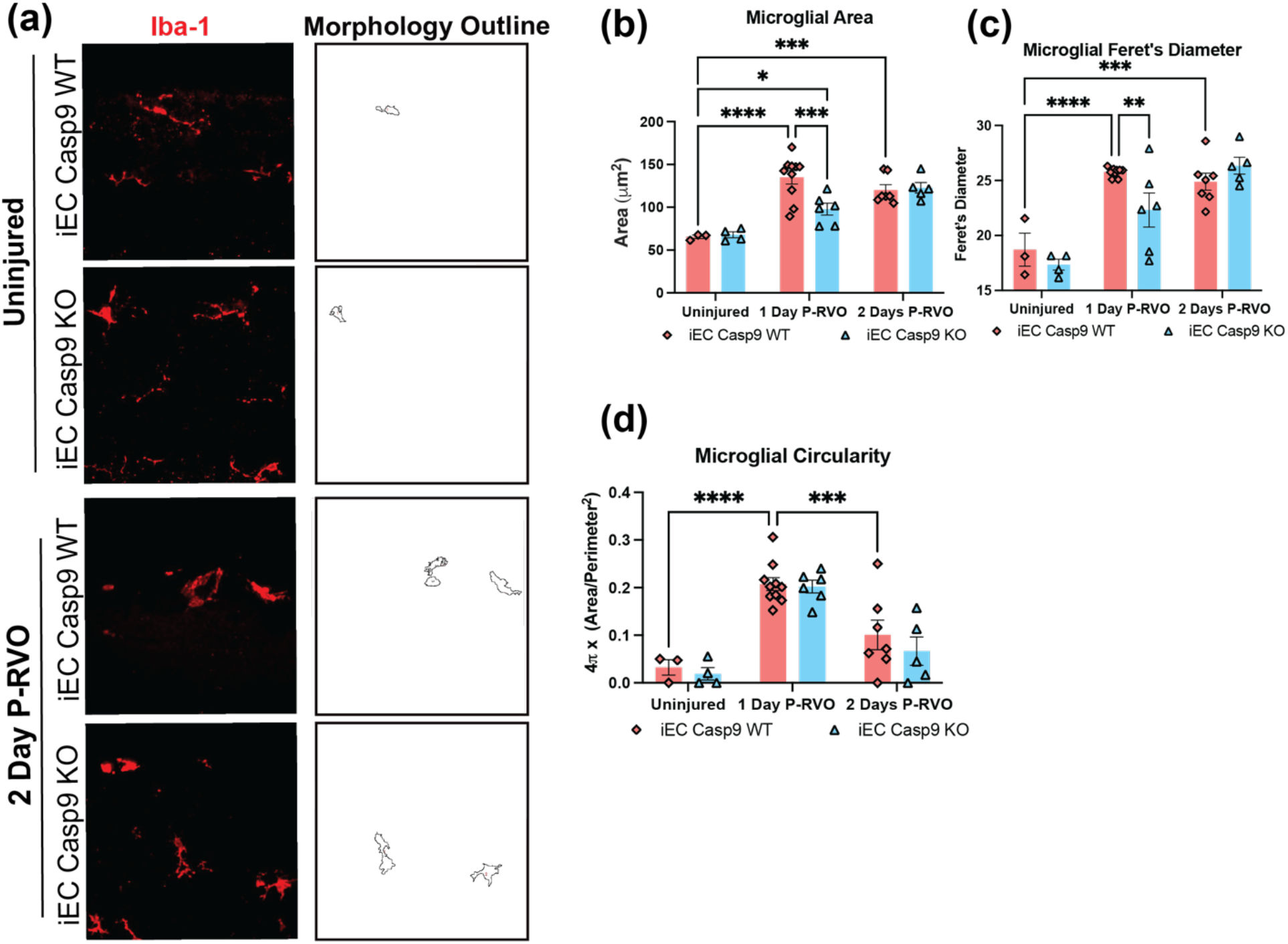
RVO induced microglial morphology changes. **(a)** Retinal cross-sections from uninjured and two-days P-RVO iEC Casp9 WT and KO mice stained with Iba-1 (red) and morphology outline (white). **(b)** Microglial area, **(c)** Ferret’s diameter, and **(d)** circularity of uninjured iEC Casp9 WT (n=5) and KO (n=6), iEC Casp9 WT (n=8) and KO (n=9) one day P-RVO, and iEC Casp9 WT (n=7) and KO (n=9) two days P-RVO. Error bars mean ± SEM; Twoway ANOVA, Fisher’s LSD test.

**Supplementary Figure 2.**
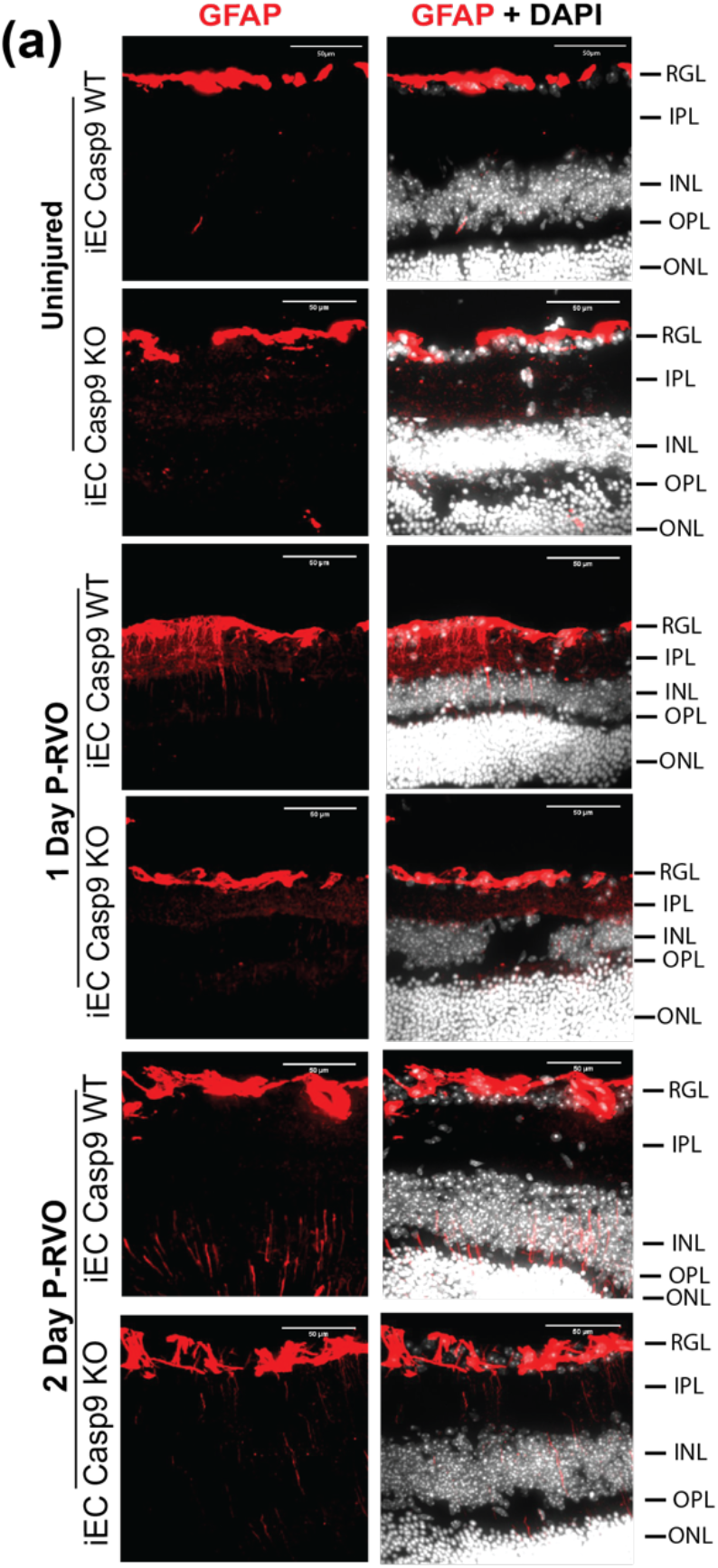
EC Casp9 promotes GFAP expression in macroglia. **(a)** Retinal cross-sections from uninjured, one and two-days P-RVO iEC Casp9 WT and KO mice stained with GFAP (red) and DAPI (white). Quantified in Figure 3e.

**Supplementary Figure 3.**
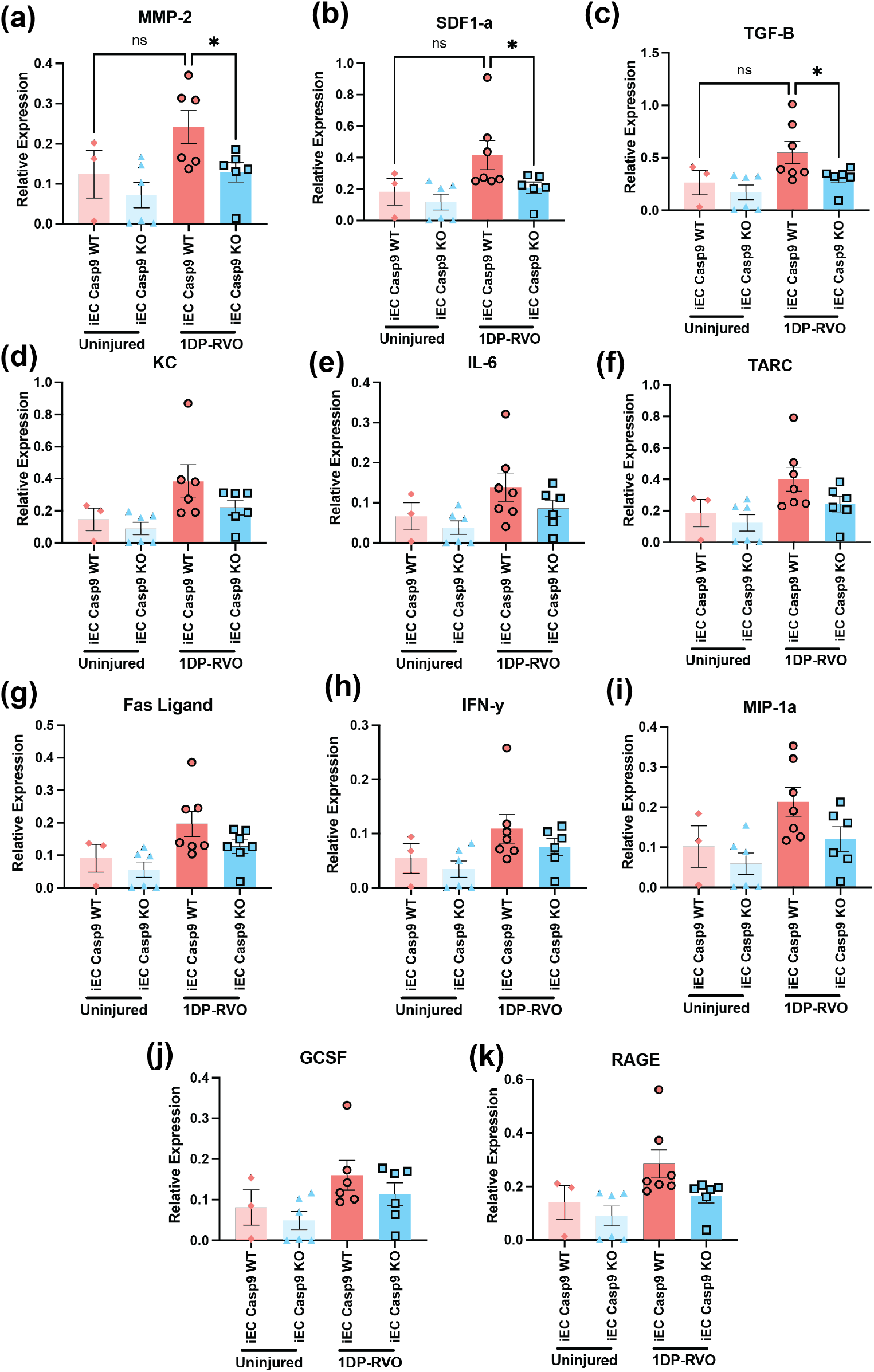
Cytokines that are not modulated by RVO. **(a-k)** Relative expression of cytokines in retinas from uninjured iEC Casp9 WT/KO and in iEC Casp9 WT/KO mice one day P-RVO.

**Supplementary Figure 4.**
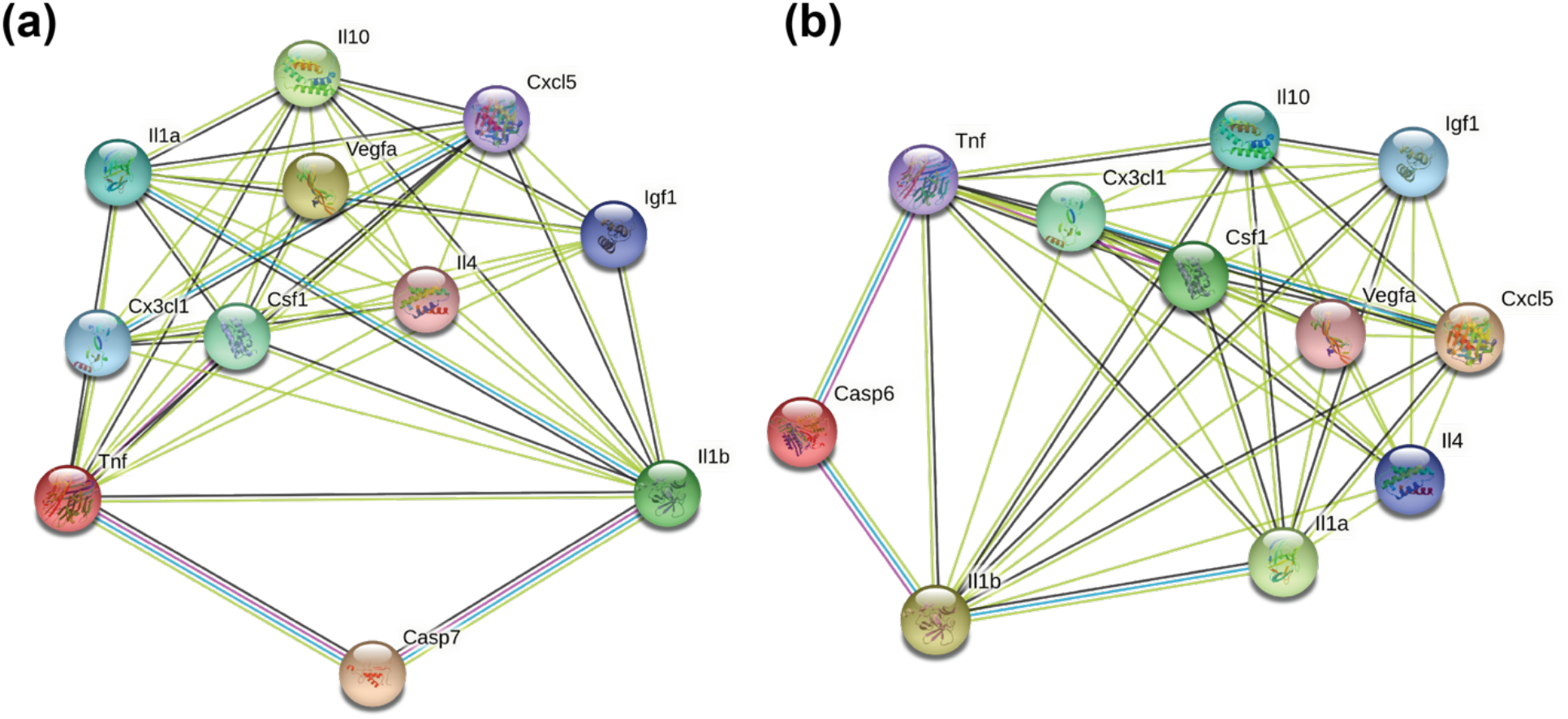
STRING analysis of EC Casp9 modulated cytokines by downstream caspase-9 targets caspase-7 and caspase-6. Protein-protein network interaction of cytokines regulated by EC Casp9 P-RVO by **(a)** caspase-7 and **(b)** caspase-6.

## Notes

### Competing Interest Statement

The authors have declared no competing interest.

